# Novel 3D Approach to Model Non-Alcoholic Fatty Liver Disease using human Pluripotent Stem Cells

**DOI:** 10.1101/2024.02.07.579290

**Authors:** Carola Maria Morell, Samantha Grace Tilson, Rute Alexandra Tomaz, Arash Shahsavari, Andi Munteanu, Giovanni Canu, Brandon Tyler Wesley, Marion Perrin, Imbisaat Geti, Subhankar Mukhopadhyay, Francesca Mazzacuva, Paul Gissen, Jose Garcia-Bernardo, Martin Bachman, Casey Allison Rimland, Fotios Sampaziotis, Irina Mohorianu, Ludovic Vallier

**Affiliations:** Wellcome-MRC Cambridge Stem Cell Institute, JCBC Cambridge Biomedical Campus, Dept of Surgery, University of Cambridge, Cambridge, UK; Liver Disease Branch, NIDDK, NIH, Bethesda, Maryland, USA; Wellcome Trust Sanger Institute, Hinxton, UK; UCL Great Ormond Street Institute of Child Health, London, London, UK; NIHR Great Ormond Street Hospital Biomedical Research Centre, London, UK; Medicines Discovery Catapult, Alderley Park, UK; Immunopathogenesis Section, Laboratory of Parasitic Diseases, NIAID, NIH, Bethesda, MD; Berlin Institute of Health Centre for Regenerative Therapies, Berlin, Germany; Max Planck Institute for Molecular Genetics, Berlin, Germany

**Keywords:** hiPSCs, disease modelling, hepatocytes, liver, NAFLD

## Abstract

**Background and aims:** Non-alcoholic fatty liver disease (NAFLD) is a major health care challenge and new therapies are urgently needed. However, the mechanisms underlying disease remain to be understood. Indeed, studying NAFLD remains challenging due to the lack of model systems recapitulating the different aspects of the human pathology. Human induced pluripotent stem cells (hiPSCs) offer a unique opportunity to address this limitation since they can be differentiated into large quantity of liver cells. Here, we took advantage of hiPSCs to develop a multi-cellular platform mimicking the complex interplays involved in NAFLD progression.

**Methods:** hiPSCs-derived hepatocyte like cells (HLCs), cholangiocytes, stellate cells, and macrophages were co-cultured in a collagen-based 3D system to reproduce the liver microenvironment. Fatty acid treatments led to a NAFLD phenotype involving cell-cell interactions which were investigated by transcriptomic and functional analyses.

**Results:** Hepatic cells were grown up to 4weeks in 3D, retaining key functions and markers. Importantly, co-cultured cells spontaneously reorganised into physiologically relevant connections: HLCs arranged around biliary structures, which established contacts with stellate cells, while macrophages organised around HLCs. Fatty acid treatments induced steatosis and lipotoxicity in HLCs. Furthermore, fat-laden HLCs prompted a non-parenchymal cells response altering tissue architecture.

**Conclusions:** Our multicellular platform provides a new approach to model interactions between human hepatic cells during NAFLD progression. Such approach has the potential to investigate the sequential events driving chronic liver diseases, including hepatocellular injury, inflammation and fibrosis. Furthermore, our system provides a unique and urgently needed tool to investigate the molecular mechanisms associated with NAFLD and ultimately to validate new targets for therapeutics development.

**List of abbreviations:** COs, cholangiocytes organoids; FFA, free fatty acids; hiPSCs, human induced pluripotent stem cells; HLCs, hepatocyte like cells; HSCs, hepatic stellate cells; M0, hiPSCs-derived macrophages; NAFLD, non-alcoholic fatty liver disease; NPCs, non-parenchymal cells; OA, oleic acid; PA, palmitic acid.

## INTRODUCTION

Non-alcoholic fatty liver disease (NAFLD) designates a group of diseases sharing a wide range of hepatic manifestations starting with hepatocytes steatosis, the first pathogenic step towards steatohepatitis and fibrosis, followed by cirrhosis and ultimately liver failure or cancer. The incidence of NAFLD is exponentially growing due to the pandemic of metabolic disorders associated with obesity. Approximately 30% of the general population could be affected by NAFLD in developed countries[1, 2]. Our understanding of the mechanisms driving NAFLD is still incomplete and there are no available therapies to slow disease progression. Currently, therapeutic interventions focus on lifestyle modifications, while organ transplantation is the only option for end stage disease, making NAFLD one of the leading causes of liver transplant[2, 3].

NAFLD is a multi-cellular disease. Chronic lipid accumulation in hepatocytes eventually leads to cellular damage. This scenario triggers an intense cross-talk between hepatic non-parenchymal cells (NPCs) (cholangiocytes, hepatic stellate cells [HSCs] and Kupffer cells), which strongly influences the pathologic phenotype and disease progression. Hepatocytes lipotoxicity provokes inflammation via macrophages activation, fibrosis orchestrated by HSCs, and ductular reaction involving cholangiocyte proliferation[4, 5]. Thus, understanding cellular interplays is essential to develop new therapies.

Animal models are commonly used to investigate the mechanisms underlying NAFLD and have provided important knowledge regarding disease pathogenesis. Accordingly, a wide range of dietary or genetic murine models are currently available to study NAFLD[6]. However, these models don’t reproduce the full spectrum of the human pathophysiology, and they are not easily compatible with large scale drug screening. Furthermore, their use is becoming increasingly ethically questionable. Thus, a major effort is ongoing to develop humanised *in vitro* platforms providing a complementary approach. A variety of *in vitro* systems has been developed[7–9], which have established a proof of principle that NAFLD can be modelled *in vitro* for testing drugs currently in clinical trials. However, they rely on primary human cells which are only available in limited supply. Thus, these *in vitro* platforms remain challenging to use for high-throughput screenings. Alternatively, transformed hepatic cell lines are often used, but their malignant background decreases their metabolic relevance[9, 10].

An advantageous alternative to primary cells is offered by human induced pluripotent stem cells (hiPSCs) since they can produce large quantities of not only hepatocytes, but a diversity of hepatic cells including macrophages[11], stellate cells[12] and cholangiocytes[13]. hiPSC-derived hepatocytes (HLCs) recapitulate key properties of their *in vivo* counterparts[14–17], can be produced from a diversity of patients, and have been proven useful to model liver disorders[18–20]. Recently, multicellular hiPSCs-based approaches have been used to model NAFLD. In one study[21], hiPSC-derived hepatoblasts were used to generate hepatic organoids comprising both hepatocytes and cholangiocytes, which allowed to model steatosis, lipotoxic damage and ductular reaction. Interestingly, lipid-related metabolic pathways were dysregulated, revealing a transcriptomics signature similar to NAFLD patients. However, this model was limited by the absence of mesenchymal cells, therefore lacking inflammatory and fibrotic components. To include NPCs, another report[22] took advantage of foregut spheroids to generate liver organoids containing mainly HLCs, followed by HSCs, while biliary-like cells and macrophages were present in small percentages. This allowed to mimic aspects of steatosis, inflammation and fibrosis in a dose and time-dependent fashion. However, in this model, the precise identity and percentages of cells could not be regulated, giving rise to heterogeneous cellular aggregates without functional organisation.

To address these limitations, we developed a novel 3D *in vitro* platform that captures hepatic multicellularity and liver organization. We recreated the hepatic microenvironment by co-culturing HLCs with NPCs in a physiological ratio using a 3D collagen matrix. Hepatic cells generated *in vitro* self-assembled to mimic the architecture of the liver and maintained key features for up to 4-weeks in culture. We used this platform to model NAFLD and we were able to recapitulate several aspects of the disease, from hepatic steatosis to NPCs response. Moreover, transcriptomic and lipidomic analyses revealed a NAFLD signature in HLCs, while NPCs displayed inflammatory and fibrotic responses. Our platform offers a new approach to model NAFLD reproducing key aspects of the disease and providing a new tool to study the human pathogenesis. Ultimately, our system will enable the discovery and validation of novel therapeutic targets.

## METHODS

*Cell treatments, Reporter cell lines, Cytochrome P450 activity, Presto Blue assay, Albumin/AFP ELISA, Bile acids mass spectrometry, Lipidomics, Quantitative Real Time PCR (qPCR), Immunofluorescence, and Flow protocols, RNA sequencing and statistical analyses, are detailed in Supplementary Information*.

### Cell Culture

Please refer to Supplementary Tables1-2 for a complete description of cell culture reagents. The hiPSC line A1ATD^R/R^[16, 19] was cultured on 10µg/ml vitronectin coated plates and maintained in chemically defined medium (in house E8) supplemented with 2ng/mL TGFβ1 and 25ng/mL FGF2, as previously reported[23]. hiPSCs were used to generate hepatocyte like cells (HLC), following our protocol[14] (summarised in supplementary table3), and to obtain macrophages (M0) as previously reported[11]. Cholangiocytes organoids (CO) were obtained from primary tissues[24], while the human cell line LX2[25] was used to model stellate cells.

### 3D (co)cultures

The 96-wells RAFT system (Lonza) was chosen for 3D cultures. At D23 of differentiation, HLCs were dissociated in small clumps with Cell Dissociation buffer, and seeded in the RAFT following the manufacturer’s instruction at a density of 70’000cells. COs were treated with Cell Recovery Solution for 30 minutes, and clumps were mechanically broken into smaller aggregates with a p1000 pipette; LX2 were detached with Trypsin-EDTA, while M0 at D7 of differentiation were collected after 10 minutes treatment with TrypLE express. NPCs were seeded in the RAFT at a density of 50’000cells for monocultures. In simple co-cultures, the ratio was 70%HLCs/30%NPCs, while in complex co-cultures ratio was 60%HLCs/5%COs/10%HSCs/25%M0. The co-culture medium (CBD medium) was supplemented with 20ng/mL OSM and 50ng/mL HGF.

### Bulk RNA-sequencing

RNA was extracted from 3 biological replicates of control, PA and OA treated (24h and 1week) samples using the RNeasy Micro Kit (Qiagen). Library preparation and sequencing was performed at the Wellcome Trust Sanger Institute (Hinxton, UK). RNA was poly-A selected and used for library preparation with the NEBNext Ultra II Directional RNA Library Prep Kit for Illumina. Sequencing was run on an Illumina HiSeq v4 to obtain 75bp paired-end reads, generating at least 100 million reads per sample. To assist with the filtering of reads, FastQC was performed on all samples to ensure that the quality threshold was above 30 throughout the entire read length. For analyses please refer to Supplementary Information. The RNA-sequencing dataset is available at Array Express under the accession number E-MTAB-10598.

### Single cell RNA-sequencing (scRNA-seq)

*Samples preparation –* RAFTs were rinsed with PBS before treatment with DNAse [200U] (D4627, Sigma)/Liberase [0.2U] (5401160001, Sigma) solution in culture medium. Samples were left on a shaker for 15 minutes at 37C to completely digest the matrix. Cells suspension was washed in warm medium+10%FBS and further dissociated into single cells via gentle pipetting and 4 minutes of StemPro™ Accutase™ incubation at 37C, followed by 2 washes in cold medium. To ensure a sufficient number of viable cells was retrieved, 5 technical replicates were pooled together. Single-cell suspension was processed into single-cell emulsions using the 10X Genomics Chromium controller. Each individual cell was encapsulated in an oil-based droplet and individual transcripts were barcoded using short DNA sequences containing both an identifier for the cell and individual transcript. A reverse transcription immediately following the formation of this emulsion converted the transcripts into stable DNA and was stored at −25C. Library preparation using V2 chemistry was performed using the recommended protocol from 10X Genomics.

*scRNA-Seq analyses:* Quality assessment of raw fastq files was performed with FastQC v0.11.3 and MultiQC v1.8. Checks included nucleotide composition and distribution of GC-content. Reads were aligned to the *H. sapiens* reference genome (GRCh38) using 10X cellranger v3.1.0; 3,449 cells were detected using default filters. The resulting count matrix was analysed in R (version 3.6.3), using custom scripts. The quality of the quantification was assessed on distributions of sequencing depth, number of unique features, and mitochondrial and ribosomal protein-coding expression levels per cell. Based on inspection of these distributions, cells with <80,000 UMIs and >1000 detected genes were retained for downstream analysis, totalling 1,519 cells. For details on normalisation, clustering and identification of marker genes, please refer to Supplementary Information. The scRNA-sequencing dataset is available under the accession number GSM4824487.

## RESULTS

### HLCs maintain key markers and functions in long term 3D cultures

To develop co-culture conditions mimicking the liver micro-environment, we took advantage of the RAFT system which allows the production of 3D collagen scaffolds. Importantly, we previously showed that this system supports the functionality of hepatocyte like cells (HLCs)[26]. To further reinforce these results, HLCs differentiated in 2D were seeded into RAFTs at a density of 70’000cells/well and grown for short term (1-week) or long term (4-weeks) studies (Fig 1A); HLCs were seeded in 3D as clumps of cells to preserve cell-cell junctions which are crucial for hepatocyte functionality and survival[26]. HLCs grown in 3D retained the expression of hepatocyte markers, such as albumin (ALB), hepatocyte nuclear factor 4 alpha (HNF4α), alpha-1-antitrypsin (protein α1AT; gene SERPINA1) and transthyretin (TTR) at similar levels compared to HLCs maintained in 2D or primary human hepatocytes (PHH)(Fig 1B). Interestingly, HLCs seemed to improve their functional maturation in 3D as shown by the increase in ALB and SERPINA1 expression over time (Fig 1B-C). HLCs grown in the RAFT continued to express α-fetoprotein (AFP) and CYP3A7 (Fig 1B); nonetheless, AFP and EPCAM expression strongly decreased with time in 3D culture (Fig 1C). Furthermore, ALB/AFP secretion ratio was significantly augmented (Fig 1D) while 3D HLCs expressed significantly higher levels of CYP3A4, although still not reaching PHH level (Fig 1E). Similarly, CYP3A4 basal activity in 2D vs 3D cells (Fig 1F) indicated that the RAFT did not alter HLC functionality. 3D culture also promoted the formation of MRP2 positive bile canaliculi (Fig 1C, Supp Fig 1A), which play a key function in toxin metabolism *in vivo*. Finally, we also detected the presence of bile acids in both 2D and 3D cultures (Fig 1G and Supp Fig 1B). Altogether, these results indicate that the RAFT system does not interfere with HLCs identity, increases specific functions and allows prolonged culture, necessary to model chronic insults *in vitro*.

**Figure 1:**
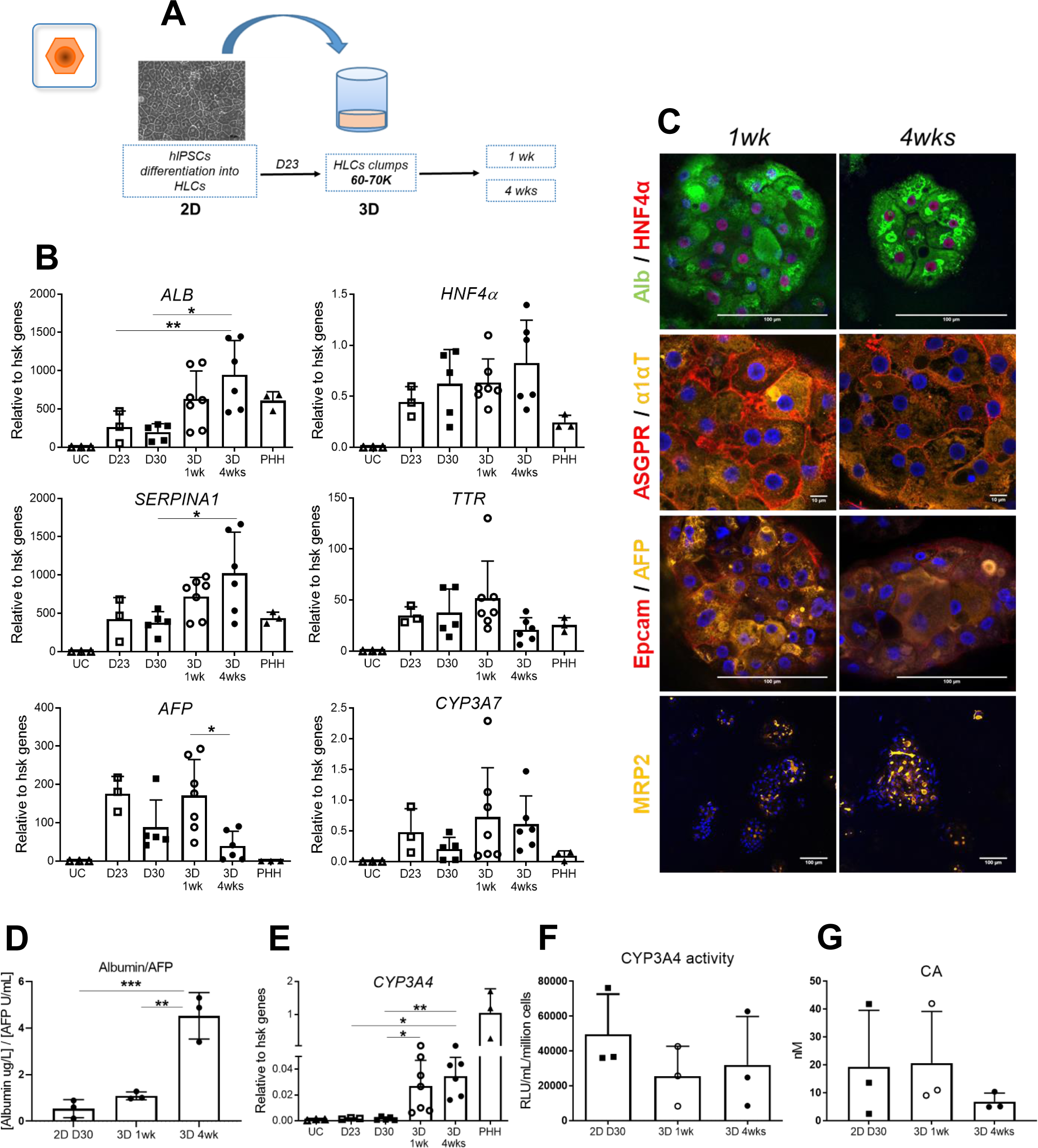
3D HLCs cultures characterisation. A) Schematic representation of HLCs embedding into RAFTs at D23. B) 2D and 3D cultures were characterised for gene expression of key hepatocyte markers (PHH and undifferentiated hiPSCs [undifferentiated control, UC] were used as positive and negative control, respectively). C) Immunofluorescence of 3D HLCs. D) ALB/AFP secretion. E) CYP3A4 expression. D) CYP3A4 basal activity. G) production of cholic acid (CA). Graphs represent average±SD of n=3-7 experiments, analysed with ANOVA.

### HLCs grown in 3D cultures allow modelling of lipid accumulation and lipotoxicity associated with NAFLD

We next decided to explore the utility of our system for modelling NAFLD. For that, HLCs were grown in 3D in the presence of oleic acid (OA) or palmitic acid (PA). OA is known to promote lipid accumulation, while PA is a saturated free fatty acid (FFA) more prone to lipotoxicity. HLCs were treated for 24hrs, 1-week or 4-weeks to model chronic injury. Exposure to OA resulted in significant lipid accumulation after 24hrs, and by 1-week lipid droplets were observed throughout HLCs aggregates (Fig 2A,B). Of note, 2D HLCs cultures failed to produce a clear steatotic phenotype (Fig 2A, top row), suggesting that the 3D environment enhanced HLCs ability to accumulate lipids. On the other hand, HLCs challenged with PA displayed limited steatosis while showing signs of cellular stress including loss of cell-cell tight junctions after 4weeks (Fig 2A, Fig 2B, Supp Fig 2A). Furthermore, exposure to PA reduced HLCs viability by 50 and 70% after 1-week and 4-weeks treatments, respectively (Fig 2D and Supp Fig 2C). We then assessed cell functionality of fat laden HLCs. Addition of PA decreased the expression of albumin suggesting that the observed cell death could be associated with dedifferentiation (Fig 2A). Furthermore, CYP3A4 basal activity was decreased by OA (4-weeks) and PA (1-week) (Fig 2C and Supp Fig 2B); this was expected as hepatocyte metabolic activity is altered in patients with NAFLD and reduced cytochrome P450 capacity has been reported[27]. These results show that our 3D platform allows the modelling of lipid accumulation and lipotoxicity induced by FFAs.

**Figure 2:**
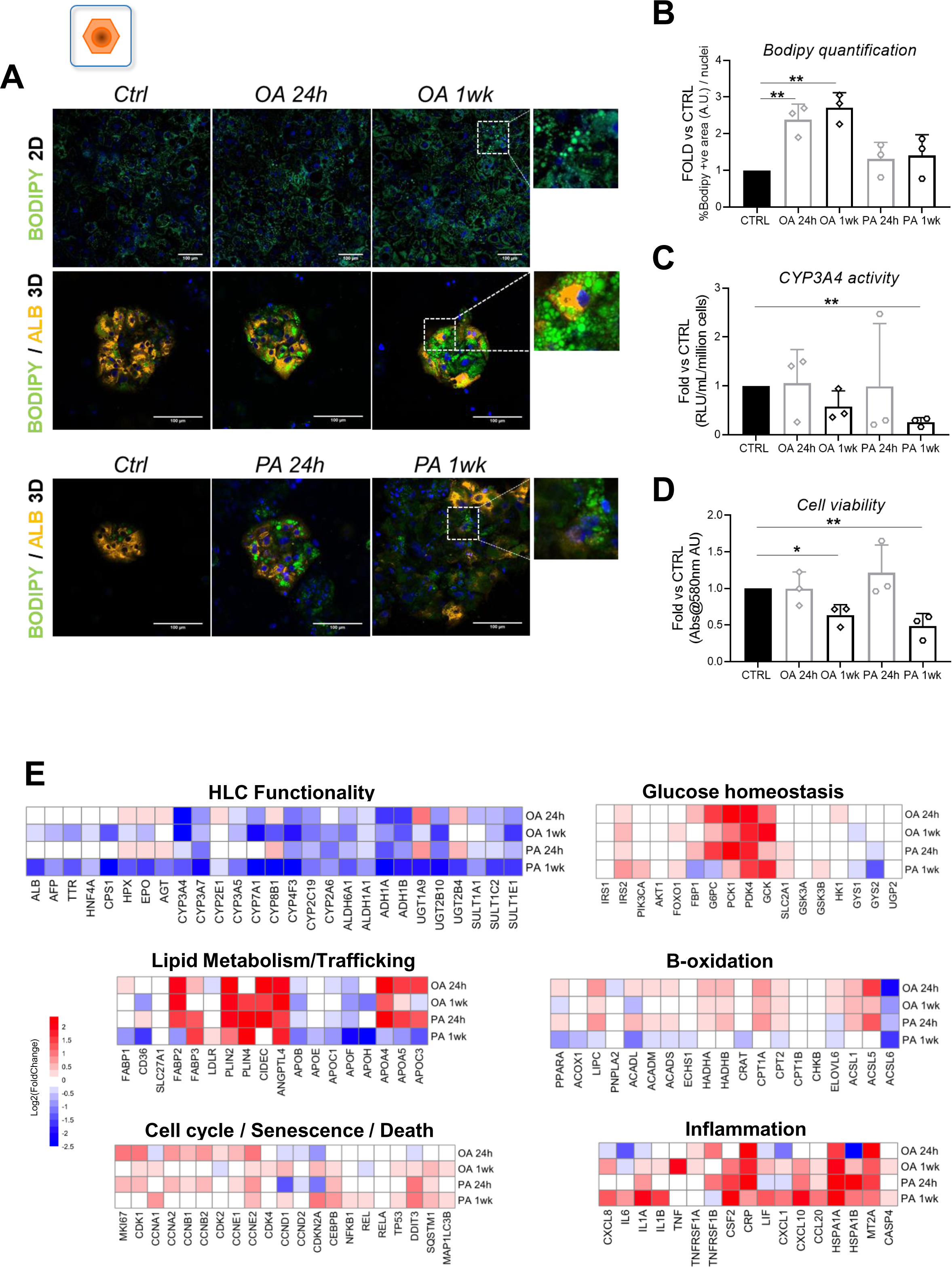
3D HLCs cultures mimic steatosis and lipotoxicity. A) FFAs challenge in HLCs visualised with Bodipy493/503 (green): top row, 2D HLCs treated with OA; middle row, OA treatment; bottom row, PA treatment. 3D cells were stained for albumin (yellow). B) Bodipy493/503 quantification, C) CYP3A4 activity and D) cell viability, all performed on 3D cultures. Graphs represent average±SD of n=3 experiments, analysed with ANOVA. E) RNA-Seq analyses on HLCs monocultures of disease-associated pathways. Heatmaps represent the log2 fold change of each treatment and timepoint vs control group. Red indicates an increase, whereas blue indicates a decrease in expression relative to control.

### Transcriptomic analyses confirm the interest of HLCs to model NAFLD

To further confirm the interest of our approach for modelling NAFLD, we performed bulk RNA-Seq analyses on HLCs exposed to FFAs for 24hrs and 1-week. Principal component analysis (PCA) showed a distinct separation of treated versus control samples depending on the length of the treatment (Supp Fig 3A) thereby confirming the impact of FFAs on HLCs. Differential gene expression analyses identified 159 downregulated and 102 upregulated genes after 24hrs treatments, while 155 and 42 genes were respectively down- and upregulated after 1-week (Supp Fig 3B-C). Gene ontology analyses revealed that 24hrs of FFAs treatment upregulated pathways related to cell cycle, lipid metabolism and transport (Supp Fig 3D). On the other hand, HLCs also downregulated pathways related to cellular adhesion, cell-cell signalling, and basic hepatocytes functions such as drug metabolism (Supp Fig 3E) and protein secretion (i.e. ALB/TTR/AFP/EPO, Fig2E). Of note, 1-week PA was the most impactful treatment for HLCs functionality. These analyses confirm that HLCs undergo steatosis and cell death associated with dedifferentiation especially when grown in the presence of PA.

We then focused on specific pathways known to be dysregulated in NAFLD (Supplemental Table4 and Fig2E). Gluconeogenesis genes (G6PC, PCK1) were upregulated, while genes associated with glycogen synthesis decreased (GYS1, GYS2). On the other hand, few insulin signalling genes (PIK3CA, IRS2, FOXO1) increased suggesting a process resembling insulin resistance. Importantly, for all groups but one (1-week PA) we registered an upregulation in genes involved in lipid uptake and trafficking, and in lipid oxidation. Interestingly, 24hrs treatments resulted in a transient increase in cell cycle-related genes (CDK1 and MKI67) suggesting that FFAs could increase proliferation in this acute phase. However, cell cycle regulators reverted to basal levels in prolonged treatments, while the expression of markers related to senescence (CDKN2A/CEBPB/TP53/NFKB), UPR (DDIT3) and autophagy (SQSTM1/MAP1LC3B) were strongly induced. Repeated exposure to FFAs upregulated pro-inflammatory genes suggesting that hepatocytes themselves could be part of the inflammation process leading to fibrosis in chronic injury. Of particular interest, CXCL8 was significantly induced by 1-week exposure to FFAs; CXCL8 is a known activator of the immune system, while significantly high levels of CXCL8 were reported in serum from NASH patients[28]. Thus, steatotic hepatocytes might play a pivotal role in NAFLD progression by promoting a pro-inflammatory environment. Taken together these results demonstrate that FFAs, especially PA, have a potent effect on HLCs functionality, metabolism and survival while inducing associated senescence, UPR and autophagy, and a strong pro-inflammatory signature.

### Lipidomic analyses reveal disease pathways in HLCs exposed to FFAs

Altered lipid metabolism is hallmark of NAFLD, and thus we decided to investigate the lipidomic profiles of our *in vitro* model. We collected supernatants after 1-week of culture in the presence of FFAs and probed the lipids secreted in each condition. PCA analyses revealed that the lipidome of OA-treated HLCs diverged strongly from control and PA-treated groups (Supp Fig 4A). This difference in lipids composition suggested that OA-treated HLCs retained specific lipid species which might result from the induction of steatosis. This is in line with the hypothesis that steatosis could be an initial response to FFAs overload. Indeed, unsaturated FFAs are more prone to be incorporated into triglycerides, and this mechanism appears to be protective against lipotoxicity; the opposite occurs with PA which, as a result, is detrimental for hepatocytes[29, 30]. These analyses revealed numerous FFAs derivatives and the main detectable products were corresponding mono and di-substituted phospholipids (phosphatidylcholines [PC] and phosphatidylethanolamines [PE]) and triglycerides (Supp Fig. 4B, Supplementary Table7). The supernatant of OA-treated HLCs was enriched with arachidonic acid, lysoPC and lysoPE (Supp Fig 4B). Interestingly, arachidonic acid is a pro-inflammatory mediator while lysoPC is linked to lipotoxicity; furthermore, several reports have described an increase of lysoPC and lysoPE in the plasma of NASH patients[31, 32]. Taken together, these data suggest that chronic OA exposure could predispose hepatocytes to lipotoxicity by releasing lipid species that support disease progression. Surprisingly, PA treatment resulted in reduced release of lysoPC, which could be explained by accumulation of these species intracellularly, but also by HLCs dedifferentiation induced by PA. In conclusion, these results reinforce the evidence that HLCs grown in the presence of FFAs acquire an injury signature related to NAFLD at the molecular level.

### Non-parenchymal cells can be grown for prolonged time in 3D while preserving their cellular identity

Several cell types play a key role in NAFLD and a multicellular platform would be useful to model the different steps characterising disease progression. Therefore, we decided to develop 3D conditions to co-culture cholangiocytes, hepatic stellate cells (HSCs) and macrophages. The first step was to address whether the 3D collagen-based environment could interfere with the activity or survival of non-parenchymal cells (NPCs). Cholangiocytes organoids (COs) (Supp Fig 5A-B) were derived from human common bile ducts[24]. COs in RAFTs organised into elongated structures with a lumen, while retaining the expression of cholangiocytes markers (Fig 3A-B) up to 4-weeks in culture.

**Figure 3:**
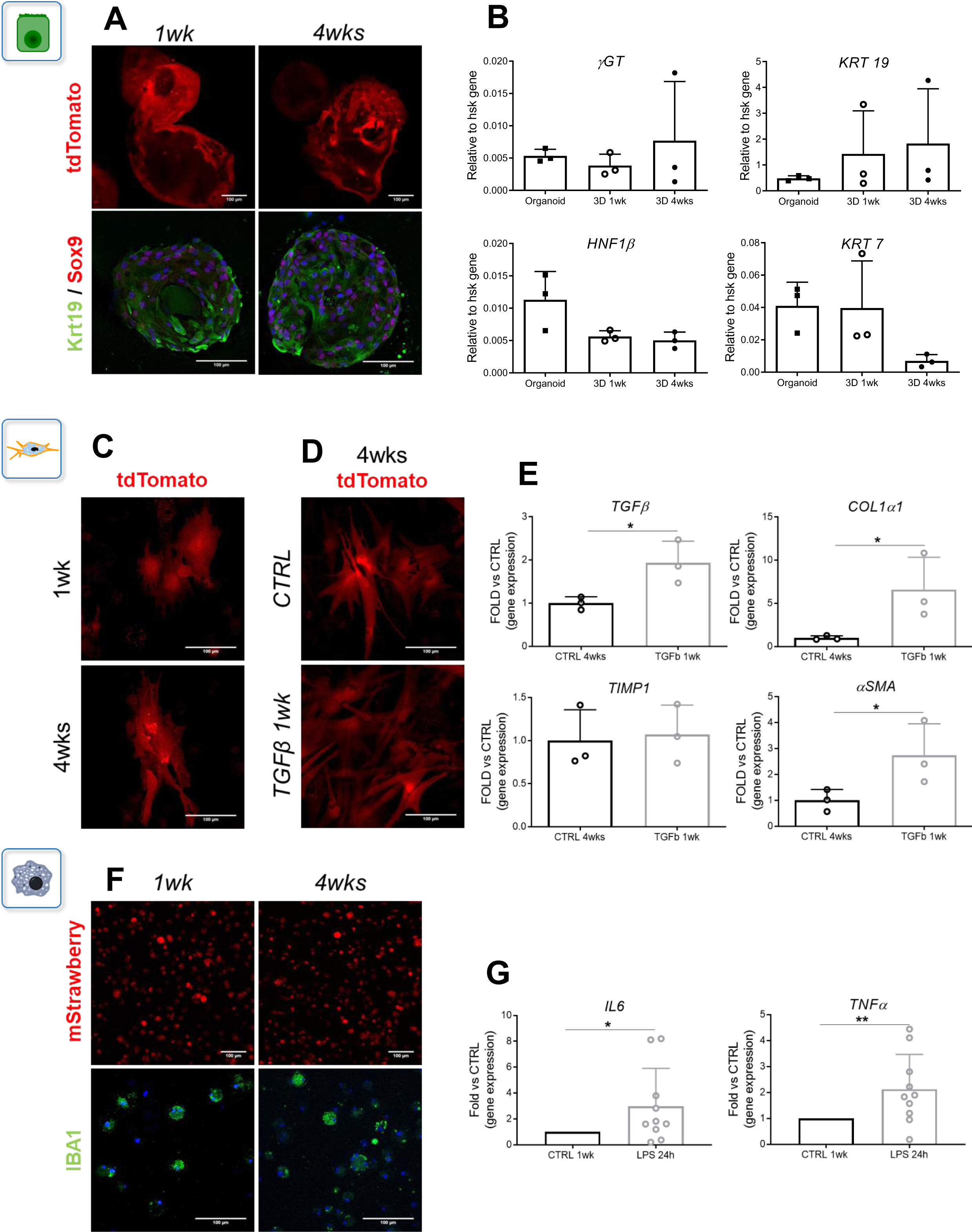
3D cultures are compatible with the growth and activation of NPCs. A) Representative pictures of COs structures in RAFTs, positive for tdTomato reporter and biliary markers; B) biliary gene expression compared to COs grown in matrigel. C) Quiescent-like morphology of HSCs-tdTomato+ grown in collagen; TGFβ1-induced D) myofibroblast-like morphology and E) pro-fibrotic genes expression. F) M0-mStrawberry+ pancake-like shape and IBA1 expression. G) Pro-inflammatory genes expression after LPS treatment. Graphs represent average±SD of n=3-10 experiments, analysed with t-Test.

The same approach was applied to HSCs taking advantage of the LX2 cell line[25]. Interestingly, HSCs in RAFTs did not display an activated morphology (Fig 3C) or phenotype as shown by the decrease in the expression of profibrotic markers TGFβ1 and collagen type I (Col1α1) (Supp Fig 6A). Other fibrotic markers were either not affected (TIMP1) or increased (αSMA) only after 4-weeks. Of note, both αSMA and GFAP (Supp Fig 6B) were expressed at basal levels in 2D or 3D, as expected with LX2 cultures. To confirm that 3D HSCs could model fibrotic response, we stimulated LX2 with TGFβ1. While efficiently promoting a profibrotic phenotype in 2D cells in 72hrs (Supp Fig 6C), a longer treatment was required to induce a fibrotic response in 3D (1-week, Supp Fig 6D-E). This activation was associated with the acquisition of a myofibroblast-like morphology (Fig 3D) and induction of fibrotic markers (Fig 3E).

Concerning Kupffer cells, we generated macrophages (M0) from hiPSCs following a well-established protocol[11]. M0 were derived in 2D and expressed typical myeloid markers CD11b and CD45 (Supp Fig 7A-C). These cells lost their phenotype after 14 days in 2D (data not shown), while M0 transferred into RAFTs retained the expression of CD14 and CD68 and did not spontaneously increase the expression of pro-inflammatory markers (Supp Fig 7D). Furthermore, 3D M0 maintained a pancake-like morphology up to 4-weeks and retained the expression of IBA1 (Fig 3F). The cells were also polarised by LPS stimulation as shown by the induction of IL6 and TNFα (Fig 3G) as reported in 2D[11] (Supp Fig 7E).

In sum, the RAFT system is suitable to grow hepatic NPCs up to 4-weeks. Each cell type maintained key phenotypical features including the capacity to react to pro-fibrotic and pro-inflammatory stimuli.

### Hepatic cells co-cultured in 3D spontaneously organize to mimic the liver microenvironment

We established conditions allowing the co-culture of these different cell types with HLCs. We started with 2 cell types at a ratio of 7:3 (HLCs:NPCs) to respect *in vivo* cell ratio. In addition, we took advantage of reporter cell lines expressing EGFP or RFP (tdTomato or mStrawberry reporters) to monitor cellular organization. Of note, a major challenge in establishing co-cultures conditions is to define media formulations compatible with all the different cell types. HLCs medium was not compatible with COs survival (data not shown). Thus, we screened several basal media and identified that William’s E supplemented with 50ng/ml HGF and 20ng/ml OSM (refer to Supplementary Tables1-2) could support all the hepatic cells included in our platform. In these conditions, HLCs rapidly surrounded the tubular-like structures formed by cholangiocytes (Supp Fig 8A, Fig 4A), while developing a well-defined canaliculi-like network which seemed to connect HLCs and biliary cells (Fig 4B). We also observed that HSCs progressively localised around HLC clusters (Supp Fig 8B, Fig 4C) without direct interactions, unless treated with TGFβ1, which induced a myofibroblast-like morphology characteristic of activated HSCs while prompting more direct contacts with HLCs (Fig 4D). Finally, M0 initially dispersed in the collagen (Supp Fig 8C) and progressively colonised the spaces between HLC clumps over time (Fig 4E). Importantly, the presence of additional cell types did not interfere with HLCs functionality including ALB/AFP production or CYP3A4 activity (Fig 4F-G and Supp Fig 8D-E). Therefore, our co-culture conditions support hepatic cell types while allowing their spontaneous organisation in 3D.

**Figure 4:**
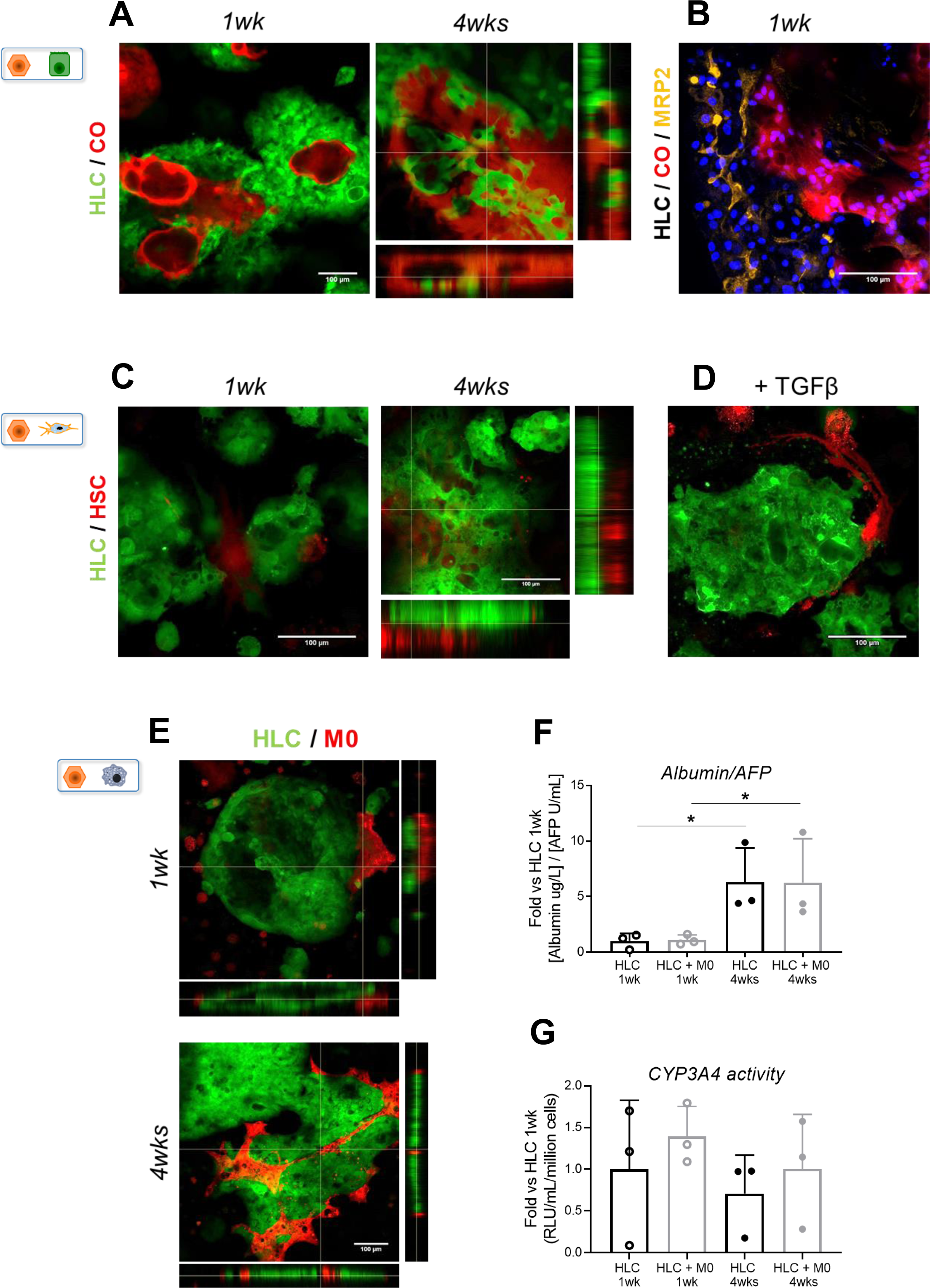
3D co-cultures promote physiologic cellular interactions. A) Live cell imaging of HLCs-EGFP+ around COs-tdTomato+, with z-stack reconstruction showing the presence of a lumen. B) MRP2 positivity of a bile canaliculi-like network after 1-week culture (HLCs-unstained, COs-tdTomato+). C) HSCs-tdTomato+ arranged on one side of HLCs-EGFP+ clumps; D) TGFβ1 treatment induced a myofibroblast-like morphology in HSCs-tdTomato+, while surrounding HLCs-EGFP+. E) M0-mStrawberry+ diffused in the RAFT, while gradually organising on one side of the HLCs-EGFP+ clumps; after 4-weeks culture, M0-mStrawberry+ spread within HLCs-EGFP+ structures. F) ALB/AFP secretion and G) basal CYP3A4 activity in 3D mono-versus co-cultures. Graphs represent average±SD of n=3-10 experiments, analysed with t-Test.

Next, we seeded the 4 cell types together at physiologic ratios: 60%HLCs/5%COs/10%HSCs/25%M0. The co-cultured cells self-organised, with HLCs surrounding COs duct-like structures within 24hrs, while HSCs and M0 dispersed around the edges of HLC clumps (Supp Fig 9A-B). Interestingly, a fraction of HSCs (Supp Fig 9C) surrounded cholangiocytes in close contact with HLCs. These interactions persisted up to 4-weeks (Fig 5A), while functionality of co-cultured HLCs was maintained as shown by CYP3A4 activity, increasing ALB/AFP secretion and bile acid production (Fig 5B-C, Supp Fig 10A-C). These results indicate that this system allows long term co-culture of hepatic cell types *in vitro* without altering HLCs functionality. Moreover, this approach enables us to precisely control the number and the cell types co-cultured, allowing the creation of a self-organised micro-environment mimicking the composition of the liver.

**Figure 5:**
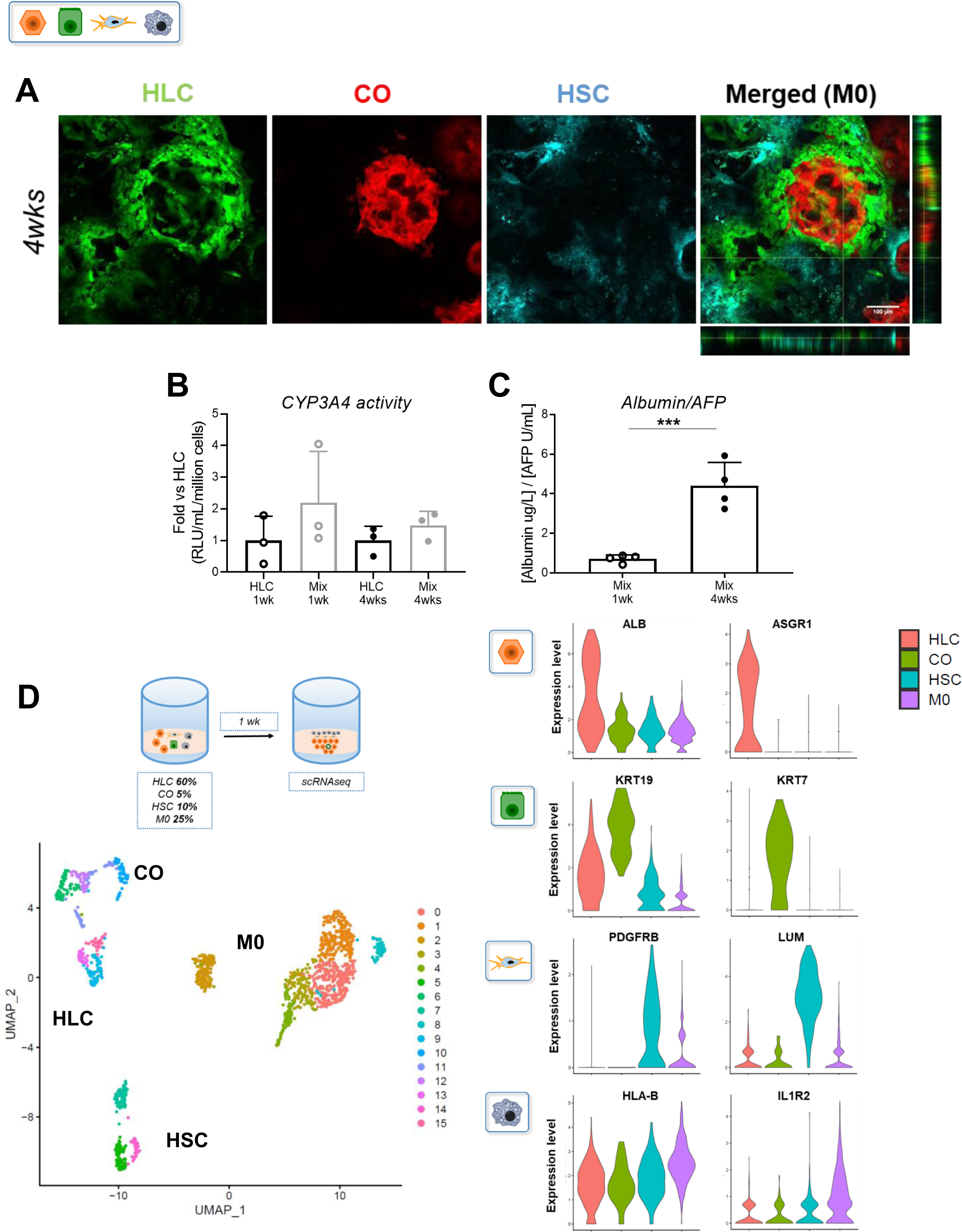
Complex 3D co-cultures mimic liver microenvironment. A) Cell-cell contacts were maintained throughout 4-weeks. HLCs-EGFP+ surrounded COs-tdTomato+, which in turn recruited HSCs-CFP+; the co-culture also included M0-unstained. B) CYP3A4 activity and C) ALB/AFP secretion in 3D mono-versus co-cultures. “Mix” indicates 4 cell types co-cultures. Graphs represent average±SD of n=3-4 experiments, analysed with t-Test. D) scRNA-Seq analyses of 1-week co-cultures. UMAP representation of 16 different groups that organised in 4 cellular clusters, identified by typical markers shown by violin plots.

### 3D co-cultures mimic cell-cell interactions observed in the liver microenvironment

To better characterise cellular interactions in our 3D co-culture system, we performed single cell RNA-Seq (scRNA-Seq) analyses on 1-week cultures. scRNA-Seq was performed to profile a total of 3449 cells, of which 44% passed QC (1519 cells; see Methods and Supplementary information). Seurat clustering was chosen for analyses due to its stability across different numbers of features, and identified 16 groups of cells (Fig 5D). The biological relevance of these clusters was assessed by enrichment analyses of marker genes that were manually curated. Using this approach, four superclusters were identified: HLCs (317 cells), COs (63 cells), HSCs (198 cells) and M0 (941 cells), thereby confirming the presence of the main cell types originally seeded into the RAFT. Of note, the numbers of retrieved cells did not mirror the seeding ratio; this difference could be explained by technical difficulties in extracting single cells from the collagen scaffold, especially hepatocytes or cholangiocytes. Nonetheless, HLCs cluster was characterised by the expression of key hepatocyte markers (ALB/ASGR1/AFP/APOB/TTR/HPX). COs expressed biliary markers (KRT19/KRT7/MUC1) and basement membrane proteins (LAMB3/LAMC2/ITGB4) among others. HSCs were identified by PDGFRB expression and by the presence of fibrosis regulators (LUM/SPARC/LOXL1/CTHRC1/COL1A1). M0 expressed markers known to regulate innate immunity (HLA-B/IL1R2/C1R/PTX3) or ECM proteins (SRGN/CCBE1) (Fig 5D and Supp Fig 11). These observations demonstrate that the identity of each cell type is preserved in our co-culture setting.

To further understand cellular interactions, we applied the CellPhone database[33] to our single-cell data set. This approach identified a diversity of known physiologic interactions (Fig 6) associated with the function of each cell type. For example, the production of plasma proteins by HLCs seemed to be supported by NPCs via the albumin-FcRn complex, which is crucial for albumin homeostasis and stabilisation[34]. On the other hand, HLCs expressed IGF2 thereby supporting cell survival. Mesenchymal cells are an important source of cytokines and growth factors in the liver. Accordingly, HSCs expressed a diversity of mitogens including EGF, FGF and HGF while the corresponding receptors appeared to be expressed by COs or HLCs. On the other hand, HLCs and COs regulated epithelial-mesenchymal cross-talk via VEGFA and PDGF signalling, acting on respective receptors on HSCs and M0. Moreover, cells-matrix interactions (integrins/ECM proteins) would suggest that these mechanisms could be responsible for guiding the 3D self-organisation of our co-culture system. Finally, our analyses also revealed the source of WNT and Notch signalling in our culture with a specific function for HSCs, which seemed to be the primary source of RSPO, WNT ligands, and TGFβ1.

**Figure 6:**
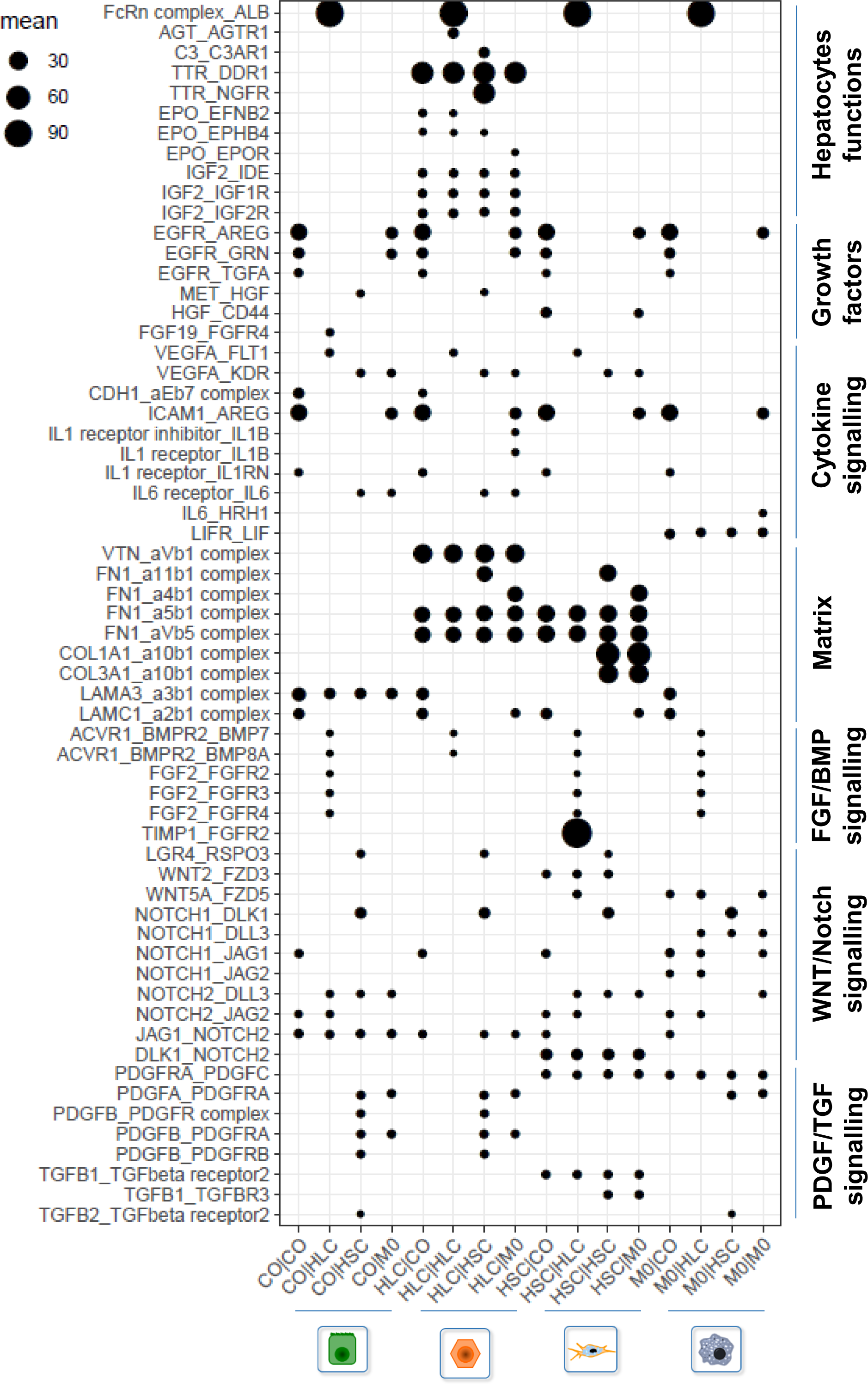
Cell-cell communication *in vitro*. CellPhone DB representation of the different cell types in 3D expressing ligands/receptors to regulate a variety of signalling. These interactions are not symmetric. The size of the dots indicates the mean-of-means (p<0.05); interactions and cell-type pairs without any dot indicate no significant interaction.

All together, these results show that our co-culture system preserves hepatic cells identity while allowing cellular interactions naturally occurring in the liver microenvironment.

### Modelling NPCs response to steatotic HLCs

The progression from steatosis to steatohepatitis is a multicellular process involving inflammation from macrophages, ductal plate reaction with cholangiocytes and fibrosis associated with HSCs. Thus, we decided to study the response of NPCs to FFAs treatment. We first evaluated the effects of FFAs on hepatic cells grown in 3D with or without HLCs. COs and HSCs were found to uptake a limited amount of lipids when grown in the presence of OA, while PA treatment had little visible effect (Supp Fig 12A-B). Interestingly, HSCs acquired an activated morphology when grown with HLCs in the presence of OA (Supp Fig 12C). FFAs had no effect on M0 in terms of lipid accumulation (Supp Fig 12D) but seemed to induce recruitment around HLCs clumps (Supp Fig 12E). We then combined the 4 cell types and challenged them with FFAs (Fig 7A). HLCs accumulated lipid droplets in the presence of OA and underwent dedifferentiation/cell death especially after long term exposure to PA. Thus, co-culture with NPCs did not affect the main phenotypes induced by different FFAs (Fig 7B-C). In addition, HSCs seemed to proliferate while adopting an activated morphology around HLCs. Pro-inflammatory and pro-fibrotic cytokines were also detected in the medium with differences for the two treatments. OA induced a stable secretion of TNFα from 1-week treatment (Fig 7D), while chronic exposure to PA induced progressive secretion of IL6 (Fig 7E). Similarly, 4-weeks treated co-cultures also produced higher although more variable levels of TGFβ1, suggesting a pro-fibrotic response (Fig 7F). Interestingly, TNFα is known to play a key role in NAFLD development and progression, and its inhibition has been shown to decrease liver injury[35]. IL6 role in NAFLD is still controversial, since it stimulates liver regeneration but also exacerbates hepatocellular damage. Both are implicated in insulin resistance and stimulate lipogenesis. However, TNFα is more linked to steatosis development and lipids/glucose metabolism while IL6 controls inflammatory response[28]. These results suggest that our platform models in part the impact of hepatic injury on NPCs. In conclusion, 3D co-cultures grown in the presence of FFAs successfully reproduced steatosis, hepatocytes lipotoxicity and key aspects of NPCs response *in vitro*.

**Figure 7:**
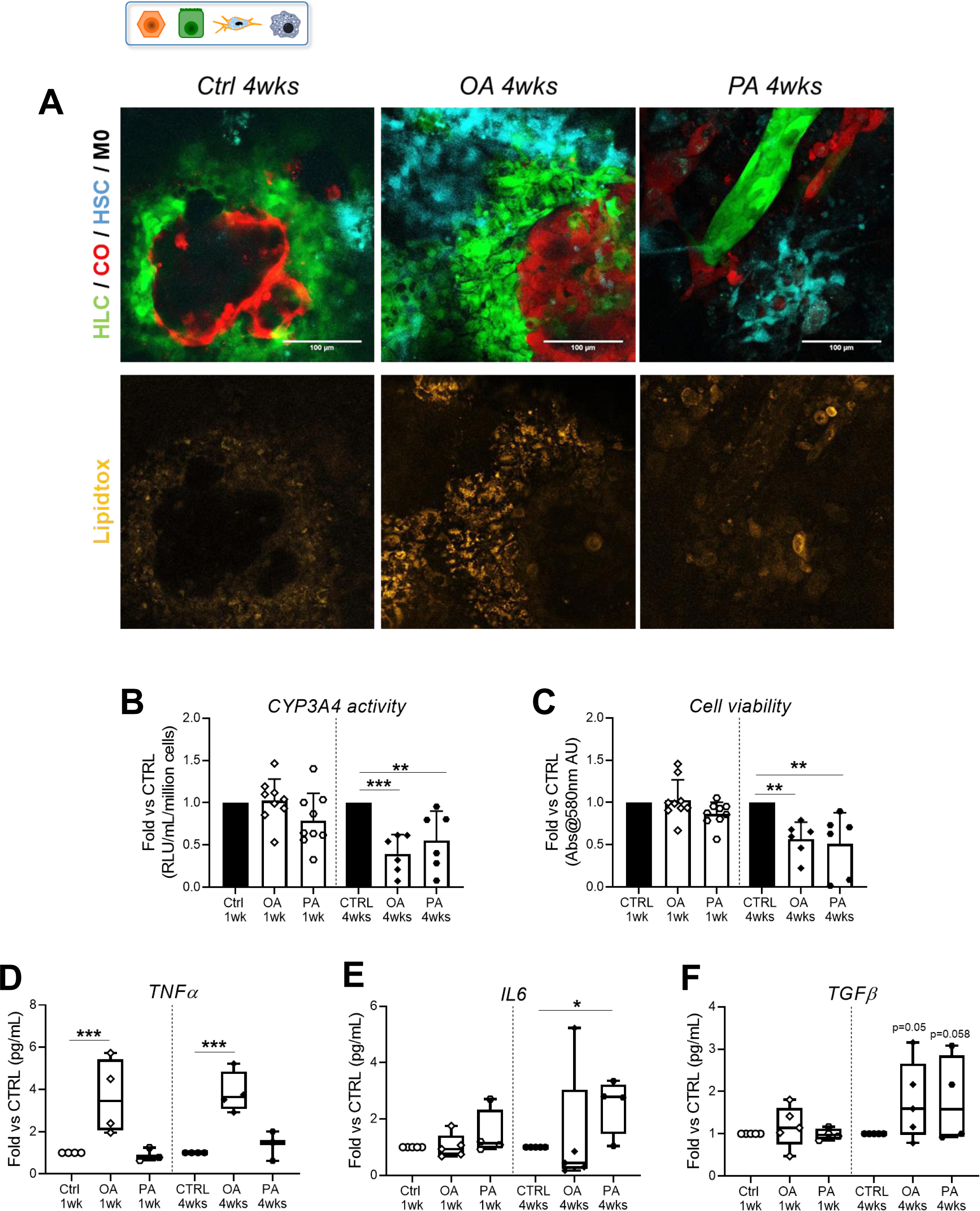
Complex 3D co-cultures promote NAFLD-like features. A) Live cell imaging shows cellular structures after long term FFAs challenge. Top row, HSCs-CFP+ accumulated around fat laden HLCs-EGFP+, that organised around COs-tdTomato+ cells after OA treatment. 4-weeks of PA resulted in the near absence of HLCs-EGFP+ clumps and unorganised COs-tdTomato+, while HSCs-CFP+ showed an activated morphology. M0s were included in the culture, but lacked a reporter gene. Bottom row, Lipidtox Deep Red (yellow) confirmed that HLCs-EGFP+ developed a steatotic phenotype after OA treatment. B) CYP3A4 activity and C) cell viability in 1-week and 4-weeks co-cultures. D) TNFα, E) IL6, and F) TGFβ1 secretion in co-cultures. Graphs represent average±SD of n=3-10 experiments, analysed with ANOVA.

## DISCUSSION

With this report, we introduce a novel multicellular hiPSCs-based hepatic platform and demonstrate its interest to model complex liver diseases, especially NAFLD. We showed how hiPSCs-derived hepatocytes (HLCs) can mimic aspects of steatosis and lipotoxicity when grown in 3D. Moreover, we were able to generate “mini livers” by combining HLCs with non-parenchymal cells (NPCs), thereby promoting cellular interactions representative of the hepatic microenvironment. This multicellular platform enabled to model hepatocytes injury observed during NAFLD progression including key associated mechanisms with inflammatory and fibrotic components.

Our approach has several advantages over currently available models. HLCs provide an unlimited source of cells which is impossible to obtain with primary cells[7, 8]. Moreover, HLCs production is robust and consistent when compared to batches of primary hepatocytes which display large variability from different donors. Furthermore, the RAFTs culture system provides a suitable environment to increase HLCs functional maturity[26]. In this study, we demonstrated that HLCs grown in 3D acquired functions which can be maintained up to 4-weeks, thereby allowing the modelling of chronic hepatic injury associated with NAFLD. Of note, HLCs have been previously used to model NAFLD. For example, lipotoxic stimuli in HLCs resulted in mitochondrial metabolism alterations[36] or interfered with lipid metabolism by inducing genes associated with disease[37]. Furthermore, our group has recently showed that HLCs provide a valuable tool to investigate the function of genetic variants associated with NAFLD, and to unravel the mechanisms by which they can influence disease progression[38]. While providing useful insights, these studies were limited to steatosis and could not address aspects linked to NPCs.

This aspect was addressed in part by using differentiation protocols resulting in heterogeneous populations of cells[21, 22]. More precisely, stochastic differentiation of foregut organoids was used to simultaneously produce several hepatic cells. However, the resulting organoids lacked functional maturation, organised 3D architecture or physiologic ratios[22]. Starting from the hepatoblast stage enabled a slightly higher control on cellular maturation, but failed to include the cells from mesoderm origin[21].

Our platform provides a solution to these limitations by precisely incorporating different hepatic cells involved in disease progression including stellate cells, macrophages and cholangiocytes. Furthermore, our approach allowed co-culture of these cell types in physiological ratios thereby mimicking the *in vivo* environment. Accordingly, combination with HLCs results in self-organisation resembling the liver architecture. scRNA-Seq analyses confirmed the functionality of these cellular interactions by revealing a diversity of interplays between these cell types. Finally, our approach allowed to control the presence of specific cell types which could provide the opportunity to investigate specific cell-cell interactions.

Importantly, our culture system could be further optimised by including endothelial cells. Indeed, it is well documented that the endothelium sustains the inflammatory and fibrotic evolution of NAFLD[39]. However, we have been so far unable to find culture conditions compatible with all the cell types included in our co-culture system and supporting endothelial cells. We used endothelial cells (HUVEC) without success, suggesting that this cell type could be particularly difficult to maintain *in vitro*.

Our approach also relies on cell types from different origins and ideally, all the cells should be generated from the same hiPSC line. This option was not implemented for the current study since production of all cell lineages from hiPSCs is complex and technically challenging even if protocols to generate cholangiocytes[13], HSCs[12], and endothelial cells[40] are now available. Nonetheless, future evaluation could involve fully derived hiPSCs platforms, which would be useful to dissect the functions of key polymorphisms without interference of genetic background.

Despite these restrictions, we showed the interest of our *in vitro* culture system to study hepatocyte injury induced by FFAs. Indeed, we demonstrated that FFAs can induce steatosis, reduce cellular function, and decrease viability to different degrees depending on the duration of treatment and the nature of the lipotoxic insult. As previously reported[30], the outcome of OA is steatosis, while PA has a lipotoxic effect, which became already evident after 1-week exposure. RNA-Seq analyses further showed that PA stimulated the expression of genes related to senescence/cell death and inflammation mediators, while affecting cellular functions. On the other hand, OA treated cells displayed an increase in the expression of genes related to lipid βoxidation. In line with these findings, OA treated cells released lipid species that were found to be increased in NAFLD patients[31, 32]. Moreover, FFAs also perturbed glucose metabolism, and the expression patterns were suggestive of insulin resistance development. Similar expression profiles were previously reported in *in vitro* models and patients’ biopsies[8], validating in part the clinical relevance of our system to model NAFLD.

Interestingly, FFAs-challenged HLCs significantly increased the expression of pro-inflammatory signals, especially CXCL8. Given the role of CXCL8 in activating the immune system, we speculated that injured HLCs could release signals creating an inflammatory environment. In addition, we also observed that TNFα was increased by OA, while PA-treated cells relied more on an IL6 response. Both cytokines are involved in NAFLD progression[28] and this could indicate a differential induction dependent on the severity of the damage or the type of injury. Intriguingly, our results showed an activation of NPCs just by enriching the medium with FFAs, without the need to add other exogenous stimuli. Taken together these results suggest that NPCs could be activated directly by injured hepatocytes and this interaction could be a relevant target for future drug development.

These data demonstrate that our platform can be used not only to model hepatocyte injury associated with NAFLD but also to uncover new mechanisms related to the disease. Thus, this novel humanised platform provides a comprehensive *in vitro* approach to identify and to validate new pathways for drug development at cell specific level.

## REFERENCES

[1] Bellentani S. The epidemiology of non-alcoholic fatty liver disease. Liver Int 2017;37 Suppl 1:81–84.

[2] Younossi ZM, Henry L. Epidemiology of non-alcoholic fatty liver disease and hepatocellular carcinoma. JHEP Rep 2021;3:100305.

[3] Konerman MA, Jones JC, Harrison SA. Pharmacotherapy for NASH: Current and emerging. J Hepatol 2018;68:362–375.

[4] Gouw AS, Clouston AD, Theise ND. Ductular reactions in human liver: diversity at the interface. Hepatology 2011;54:1853–1863.

[5] Schwabe RF, Tabas I, Pajvani UB. Mechanisms of Fibrosis Development in Nonalcoholic Steatohepatitis. Gastroenterology 2020;158:1913–1928.

[6] Ibrahim SH, Hirsova P, Malhi H, Gores GJ. Animal Models of Nonalcoholic Steatohepatitis: Eat, Delete, and Inflame. Dig Dis Sci 2016;61:1325–1336.

[7] Duriez M, Jacquet A, Hoet L, Roche S, Bock MD, Rocher C, et al. A 3D Human Liver Model of Nonalcoholic Steatohepatitis. J Clin Transl Hepatol 2020;8:359–370.

[8] Feaver RE, Cole BK, Lawson MJ, Hoang SA, Marukian S, Blackman BR, et al. Development of an in vitro human liver system for interrogating nonalcoholic steatohepatitis. JCI Insight 2016;1:e90954.

[9] Müller FA, Sturla SJ. Human in vitro models of nonalcoholic fatty liver disease. Current Opinion in Toxicology 2019;16:9–16.

[10] Godoy P, Hewitt NJ, Albrecht U, Andersen ME, Ansari N, Bhattacharya S, et al. Recent advances in 2D and 3D in vitro systems using primary hepatocytes, alternative hepatocyte sources and non-parenchymal liver cells and their use in investigating mechanisms of hepatotoxicity, cell signaling and ADME. Arch Toxicol 2013;87:1315–1530.

[11] Alasoo K, Martinez FO, Hale C, Gordon S, Powrie F, Dougan G, et al. Transcriptional profiling of macrophages derived from monocytes and iPS cells identifies a conserved response to LPS and novel alternative transcription. Sci Rep 2015;5:12524.

[12] Coll M, Perea L, Boon R, Leite SB, Vallverdu J, Mannaerts I, et al. Generation of Hepatic Stellate Cells from Human Pluripotent Stem Cells Enables In Vitro Modeling of Liver Fibrosis. Cell Stem Cell 2018;23:101–113 e107.

[13] Sampaziotis F, de Brito MC, Madrigal P, Bertero A, Saeb-Parsy K, Soares FAC, et al. Cholangiocytes derived from human induced pluripotent stem cells for disease modeling and drug validation. Nat Biotechnol 2015;33:845–852.

[14] Hannan NR, Segeritz CP, Touboul T, Vallier L. Production of hepatocyte-like cells from human pluripotent stem cells. Nat Protoc 2013;8:430–437.

[15] Hay DC, Zhao D, Fletcher J, Hewitt ZA, McLean D, Urruticoechea-Uriguen A, et al. Efficient differentiation of hepatocytes from human embryonic stem cells exhibiting markers recapitulating liver development in vivo. Stem Cells 2008;26:894–902.

[16] Rashid ST, Corbineau S, Hannan N, Marciniak SJ, Miranda E, Alexander G, et al. Modeling inherited metabolic disorders of the liver using human induced pluripotent stem cells. J Clin Invest 2010;120:3127–3136.

[17] Segeritz CP, Rashid ST, de Brito MC, Serra MP, Ordonez A, Morell CM, et al. hiPSC hepatocyte model demonstrates the role of unfolded protein response and inflammatory networks in alpha1-antitrypsin deficiency. J Hepatol 2018;69:851–860.

[18] Cayo MA, Cai J, DeLaForest A, Noto FK, Nagaoka M, Clark BS, et al. JD induced pluripotent stem cell-derived hepatocytes faithfully recapitulate the pathophysiology of familial hypercholesterolemia. Hepatology 2012;56:2163–2171.

[19] Yusa K, Rashid ST, Strick-Marchand H, Varela I, Liu PQ, Paschon DE, et al. Targeted gene correction of alpha1-antitrypsin deficiency in induced pluripotent stem cells. Nature 2011;478:391–394.

[20] Zhang S, Chen S, Li W, Guo X, Zhao P, Xu J, et al. Rescue of ATP7B function in hepatocyte-like cells from Wilson’s disease induced pluripotent stem cells using gene therapy or the chaperone drug curcumin. Hum Mol Genet 2011;20:3176–3187.

[21] Ramli MNB, Lim YS, Koe CT, Demircioglu D, Tng W, Gonzales KAU, et al. Human Pluripotent Stem Cell-Derived Organoids as Models of Liver Disease. Gastroenterology 2020;159:1471–1486 e1412.

[22] Ouchi R, Togo S, Kimura M, Shinozawa T, Koido M, Koike H, et al. Modeling Steatohepatitis in Humans with Pluripotent Stem Cell-Derived Organoids. Cell Metab 2019;30:374–384 e376.

[23] Chen G, Gulbranson DR, Hou Z, Bolin JM, Ruotti V, Probasco MD, et al. Chemically defined conditions for human iPSC derivation and culture. Nat Methods 2011;8:424–429.

[24] Sampaziotis F, Justin AW, Tysoe OC, Sawiak S, Godfrey EM, Upponi SS, et al. Reconstruction of the mouse extrahepatic biliary tree using primary human extrahepatic cholangiocyte organoids. Nat Med 2017;23:954–963.

[25] Xu L, Hui AY, Albanis E, Arthur MJ, O’Byrne SM, Blaner WS, et al. Human hepatic stellate cell lines, LX-1 and LX-2: new tools for analysis of hepatic fibrosis. Gut 2005;54:142-151.

[26] Gieseck RL, 3rd, Hannan NR, Bort R, Hanley NA, Drake RA, Cameron GW, et al. Maturation of induced pluripotent stem cell derived hepatocytes by 3D-culture. PLoS One 2014;9:e86372.

[27] Fisher CD, Lickteig AJ, Augustine LM, Ranger-Moore J, Jackson JP, Ferguson SS, et al. Hepatic cytochrome P450 enzyme alterations in humans with progressive stages of nonalcoholic fatty liver disease. Drug Metab Dispos 2009;37:2087–2094.

[28] Braunersreuther V, Viviani GL, Mach F, Montecucco F. Role of cytokines and chemokines in non-alcoholic fatty liver disease. World J Gastroenterol 2012;18:727–735.

[29] McClain CJ, Barve S, Deaciuc I. Good fat/bad fat. Hepatology 2007;45:1343–1346.

[30] Ricchi M, Odoardi MR, Carulli L, Anzivino C, Ballestri S, Pinetti A, et al. Differential effect of oleic and palmitic acid on lipid accumulation and apoptosis in cultured hepatocytes. J Gastroenterol Hepatol 2009;24:830–840.

[31] Hu C, Wang T, Zhuang X, Sun Q, Wang X, Lin H, et al. Metabolic analysis of early nonalcoholic fatty liver disease in humans using liquid chromatography-mass spectrometry. J Transl Med 2021;19:152.

[32] Puri P, Baillie RA, Wiest MM, Mirshahi F, Choudhury J, Cheung O, et al. A lipidomic analysis of nonalcoholic fatty liver disease. Hepatology 2007;46:1081–1090.

[33] Efremova M, Vento-Tormo M, Teichmann SA, Vento-Tormo R. CellPhoneDB: inferring cell-cell communication from combined expression of multi-subunit ligand-receptor complexes. Nat Protoc 2020;15:1484–1506.

[34] Chaudhury C, Mehnaz S, Robinson JM, Hayton WL, Pearl DK, Roopenian DC, et al. The major histocompatibility complex-related Fc receptor for IgG (FcRn) binds albumin and prolongs its lifespan. J Exp Med 2003;197:315–322.

[35] Wandrer F, Liebig S, Marhenke S, Vogel A, John K, Manns MP, et al. TNF-Receptor-1 inhibition reduces liver steatosis, hepatocellular injury and fibrosis in NAFLD mice. Cell Death Dis 2020;11:212.

[36] Sinton MC, Meseguer-Ripolles J, Lucendo-Villarin B, Wernig-Zorc S, Thomson JP, Carter RN, et al. A human pluripotent stem cell model for the analysis of metabolic dysfunction in hepatic steatosis. iScience 2021;24:101931.

[37] Graffmann N, Ring S, Kawala MA, Wruck W, Ncube A, Trompeter HI, et al. Modeling Nonalcoholic Fatty Liver Disease with Human Pluripotent Stem Cell-Derived Immature Hepatocyte-Like Cells Reveals Activation of PLIN2 and Confirms Regulatory Functions of Peroxisome Proliferator-Activated Receptor Alpha. Stem Cells Dev 2016;25:1119–1133.

[38] Tilson SG, Morell CM, Lenaerts AS, Park SB, Hu Z, Jenkins B, et al. Modeling PNPLA3-Associated NAFLD Using Human-Induced Pluripotent Stem Cells. Hepatology 2021;74:2998–3017.

[39] Hammoutene A, Rautou PE. Role of liver sinusoidal endothelial cells in non-alcoholic fatty liver disease. J Hepatol 2019;70:1278–1291.

[40] Patsch C, Challet-Meylan L, Thoma EC, Urich E, Heckel T, O’Sullivan JF, et al. Generation of vascular endothelial and smooth muscle cells from human pluripotent stem cells. Nat Cell Biol 2015;17:994–1003.

